# Eroding species barriers: hybridising canids remain distinct around centromeres

**DOI:** 10.1101/2025.03.26.645431

**Authors:** Filip Jagoš, Stuart J.E. Baird, Markéta Harazim, Natália Martínková

**Affiliations:** Institute of Vertebrate Biology, Czech Academy of Sciences, Kvěná 8, 60300 Brno, Czechia; RECETOX, Faculty of Science, Masaryk University, Kotlářská 2, 61137 Brno, Czechia

**Keywords:** genome polarisation, introgression, geneflow, hybridisation, wolf-dog hybridisation, wolf-coyote hybridisation, wolfdog, hybrid dog breeds

## Abstract

Species barriers are shaped by variation in recombination coupled with natural selection [1–3]. Canids lost the *Prdm9* gene 40–60 million years ago [4–6], stabilising low recombination at the centromeric ends [7] of acrocentric autosomes despite overall high recombination rates [6]. We predict canid recombination architecture will tend to maintain barriers to gene flow at one end of autosomes while eroding them at the other. Polarization [8] of 31 million single nucleotide variants (SNVs) across 980 canid genomes reveals elevated barriers to gene flow at pericentromeric regions between grey wolves, coyotes and golden jackals. Highly diagnostic SNVs are six-fold enriched in pericentromeric regions. The exception is the barrier to gene flow between dogs and wolves, which accumulated divergence differently from other comparisons: genomes of domesticated dogs preserve ancient genetic diversity lost in contemporary grey wolf populations, positioning them as reservoirs of ancestral alleles. Our results show that genome architecture modulates permeability of barriers to gene flow, pointing to pericentromeric regions as targets for reinforcement selection that may maintain canid taxa.

---

Hybridisation that results in fertile offspring followed by backcrosses to a parental taxon can provide a rapid evolutionary pathway for capturing pre-adapted traits. This process is widespread across many species, including those of conservation concern, where hybridisation is often viewed as detrimental [9–12]. Critics of granting legal protection to individuals with admixed genomes under the parental taxon’s status argue that conservation efforts should focus on the parental taxon. They advocate for the prevention of breeding involving hybrids to maintain the genetic integrity of the original species. This position is appealing to some stakeholders, as it can justify the culling of hybrids, which are often perceived as threats to livestock or as obstacles to tourism.

The grey wolf (*Canis lupus*) is at the centre of conservation debates across much of its range, where conflicts often arise over whether to protect wolves or their wild and domesticated prey. A fact that complicates the argument in favour of conservation of wolves is that the wolves hybridise with other canid taxa across their distribution range, producing fertile offspring that can backcross into the wolf gene pool [11, 13–15]. There are three main axes affecting canid gene pools across the world. In the Nearctic, extensive admixture has occurred between wolves and coyotes (*Canis latrans*), leading to the recognition of the red wolf (*Canis rufus*) as a taxon of hybrid origin [13, 16–18]. In the Palearctic, free-ranging dogs, also known as village or feral dogs, have interbred with wolves, resulting in gene flow between their genomes [10, 19–21]. Additionally, dogs have been purposefully bred with wolves late after dog domestication to produce hybrid dog breeds. The hybrid dog breeds, such as the Czechoslovakian Wolfdog or the Saarloos Wolfdog, originated from crossing established dog breeds with several wolf founders, followed by backcrossing to the dogs [22–24].

Canid hybridisation has facilitated adaptive introgression, where beneficial genes are transferred between species [8]. For example, genomic regions in wolves from Western Europe that likely originated from hybridisation with dogs carry genes under positive selection related to neurotransmission and neural system development [21]. In North America, melanism in grey wolves is believed to have originated from hybridisation with dogs shortly after introduction of dogs to the Nearctic [25, 26] [but see 27], but has since spread, following the pattern predicted by Gloger’s rule [28], and has been linked to resistance to canine distemper [29]. This suggests that hybridisation, even when seen as controversial, can play a role in enhancing the adaptive potential of species, particularly in changing environments.

Here, we analyse genome-wide diversity in canids using data from whole-genome sequencing, to quantify gene flow between wolves and other canid taxa (Figure 1a). We test the hypotheses that:

1. The species barriers between wolves, red wolves, coyotes and jackals are semi-permeable and the hybridisation has been a historical reality.
2. Introgressions are detectable across species barriers.
3. Unrecognised wolf-dog hybrids can be identified in public databases within the large global sample.

**Figure 1:**
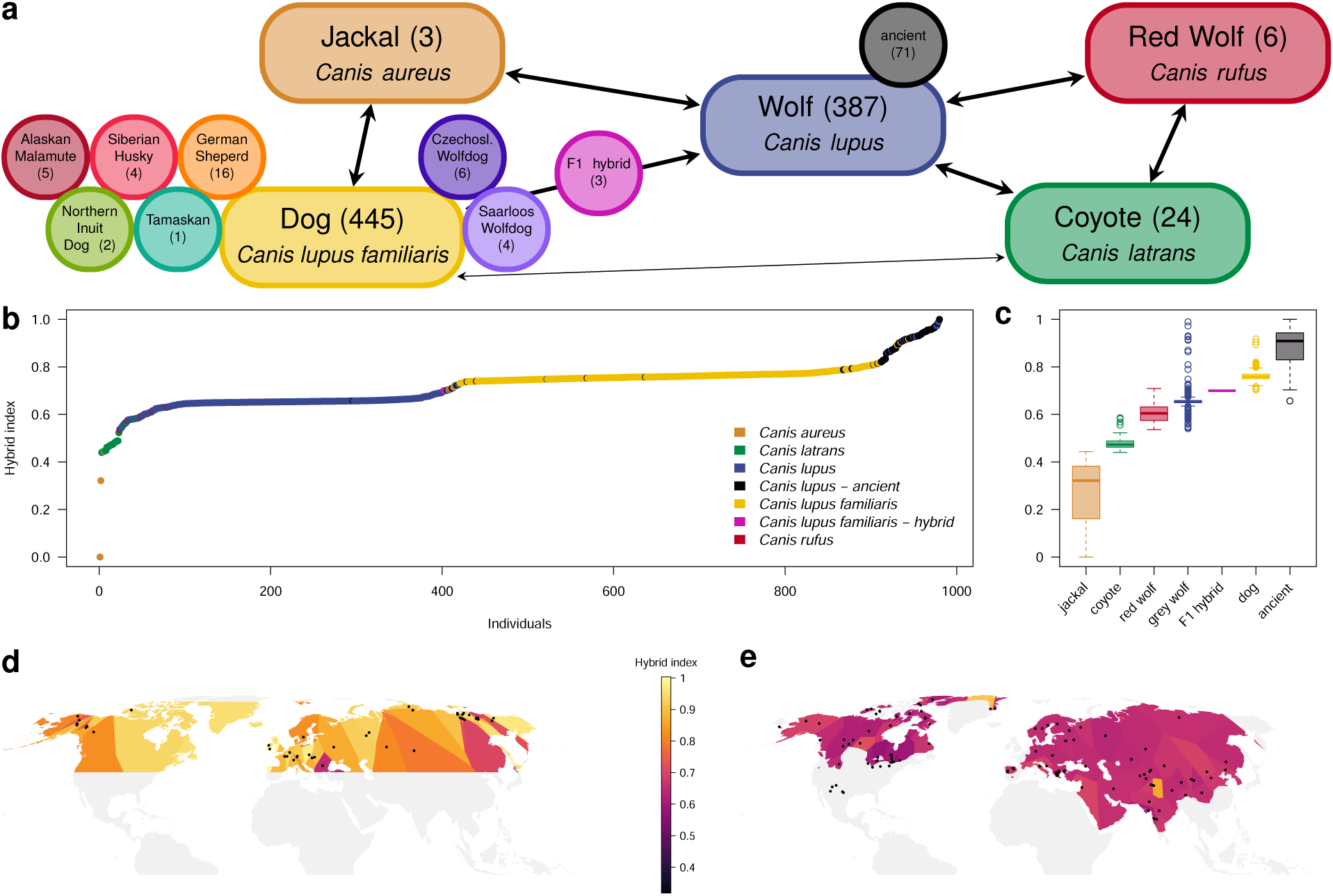
Canid admixture from polarised single nucleotide variant sites. **a**, Colour scheme of the sampling design (number of unique samples per each group in brackets). Arrows show gene flow investigated in this study. **b**, Rescaled hybrid index of 980 canid genomes shows genome admixture from 31,483,813 biallelic variant sites. Group labels were identified after genome polarisation from sample metadata (Supplementary Table 1). Dogs retain more genetic diversity present in ancient wolves than modern wolves, which have differentiated since the split from the dog ancestors. **c**, Boxplot of rescaled hybrid indices displayed in panel b. **d**, Voronoi tesselation of the rescaled hybrid indices of the ancient (*>* 1000 years old) wolf genomes with dots representing sampling sites. Tesselation was extended to 10% over the data. **e**, Voronoi tesselation of the rescaled hybrid indices of the modern wolf genomes clipped to the current grey wolf distribution range [30], showing retention of ancient genomic diversity in wolves from New Mexico, Greenland and the Tibetan Plateau. Panels d and e share the same scale of the rescaled hybrid indices.

## Canid genome admixture

Polarisation of 31,483,813 biallelic variant sites in 980 canid genomes revealed a significant axis of genetic differentiation between the golden jackal (*Canis aureus*) from Syria and a wolf subspecies, the Mexican wolf (*Canis lupus baileyi* ), from New Mexico (Figure 1b, Supplementary Table 1). Most contemporary wolves cluster closely together, adjacent to coyotes (*Canis latrans*) and red wolves (*Canis rufus*), with domesticated dogs (*Canis lupus familiaris*) positioned between these groups. Dogs are found close to Paleolithic and Neolithic grey wolves and the Mexican wolves. The sorted hybrid indices show three steep sections, indicating rapid genetic change with reduced gene flow. Following the increase in hybrid index from low values to high (Figure 1b), the steep sections are hybridisation of North American canids involving coyotes, wolves and their hybrid taxon, the red wolf, hybridisation between wolves and dogs, and historical changes in genetic diversity of wolves. We will first focus on historical changes in the wolf genetic diversity and demonstrate the retention of ancestral wolf diversity in modern dogs.

The hybrid indices of ancient wolves overlap with wolves and dogs, with dogs being overall more closely related to, albeit statistically distinct from the ancient wolves (pairwise *t*-test with FDR correction: *p*_FDR_ *<* 0.001; Figure 1c). Geographic predictions using Voronoi tessellation indicate that the genetic diversity observed in ancient wolves (Figure 1d) has largely disappeared across the wolf’s current range, with remnants persisting only in New Mexico, on Greenland, and in the Tibetan Plateau (Figure 1e). Dogs with the highest proportion of their genome similar to that of the ancient wolves are predominantly free-ranging dogs from Tibet and China (Supplementary Table 1).

When the dataset was restricted to current and ancient grey wolves and dogs from Eurasia that included 13,982,138 variant sites, the primary barrier to gene flow emerged between current and ancient grey wolves (Supplementary Figure 1, Supplementary Table 1, subset EurasiaAncient). The iris plot of the polarised genomes demonstrated that domesticated dogs have retained genetic diversity from ancient wolves, much of which has been lost from the modern wolf gene pool (Supplementary Figure 1).

## Nearctic canid hybridisation

The Nearctic canid hybridisation conundrum presents significant conservation challenges, driven by a complex interplay of historical, ecological, and genetic factors. Over the past century, human activities such as deforestation and the extirpation of wolves have facilitated the dramatic range expansion of coyotes. Once confined to the open landscapes of the southwestern Nearctic, coyotes have spread across most of the North American continent, reaching as far south as Panama and as far east as the Atlantic coast [31]. This human-mediated expansion has intensified interactions between coyotes and wolves, leading to widespread hybridisation, particularly in the eastern parts of their range. In these regions, genes introgressed from wolves into coyote populations, especially those controlling body size, are under selection, resulting in the emergence of larger coyotes in the east [13, 32]. This hybridisation has not only influenced the physical traits of eastern coyotes but has also broadened their ecological niche, allowing them to adapt to a wider range of environments [33]. In areas like the southeastern United States and the Great Lakes, the wolf-red wolf-coyote hybridisation events have created populations with mixed genetic backgrounds, complicating efforts to conserve pure species and challenging traditional species boundaries [13, 14, 16]. The endangered red wolf exemplifies the difficulties in managing and protecting hybrid populations [34, 35]. While the red wolf is an accepted species of hybrid origin between Pleistocene wolves and coyotes, recent hybridisation with a parental taxon raises concerns about genetic swamping. Conservation strategies must address the challenge of distinguishing between pure and hybrid individuals, addressing complex legal and ethical questions about their protection [12]. This requires ongoing research and adaptive management to safeguard the genetic diversity and ecological roles of the Nearctic canids.

The barrier focusing on Nearctic canids showed a gradual increase in admixture among tested samples (Figure 1b). Subsetting the data to 107 individuals of *C. latrans*, *C. rufus* and *C. lupus* from the Nearctic (Supplementary Table 1, subset NAwild) resulted in a matrix with 8,643,318 variant sites. Following genome polarisation, the hybrid index of Nearctic canids exhibited taxon-specific steps indicative of limited gene flow (Supplementary Figure 2). Using NCBI taxon labels post-hoc, four coyotes and seven wolves grouped with red wolves, whereas one red wolf was included in a group of wolves.

Most chromosomes contained large, up to 10 Mb long, heterozygous blocks in coyotes, red wolves and wolves indicative of introgressions (Figure 2a). The direction of the introgression is mainly from coyote into wolf genome.

## Wolf-dog hybridisation in the Palearctic

Although wolves coexist with other canids like jackals in the Palearctic, interspecific hybridisation has not been a significant concern. Instead, the primary admixture threat comes from domestic dogs. Historically, wolves in Europe were nearly driven to extinction, but recent decades have seen a gradual recovery and recolonisation of their former range [36–38]. While this resurgence increased connectivity amongst the distribution fragments [37, 38], the history of low numbers of wolves has brought new challenges, particularly the risk of genetic swamping from domestic dogs.

**Figure 2:**
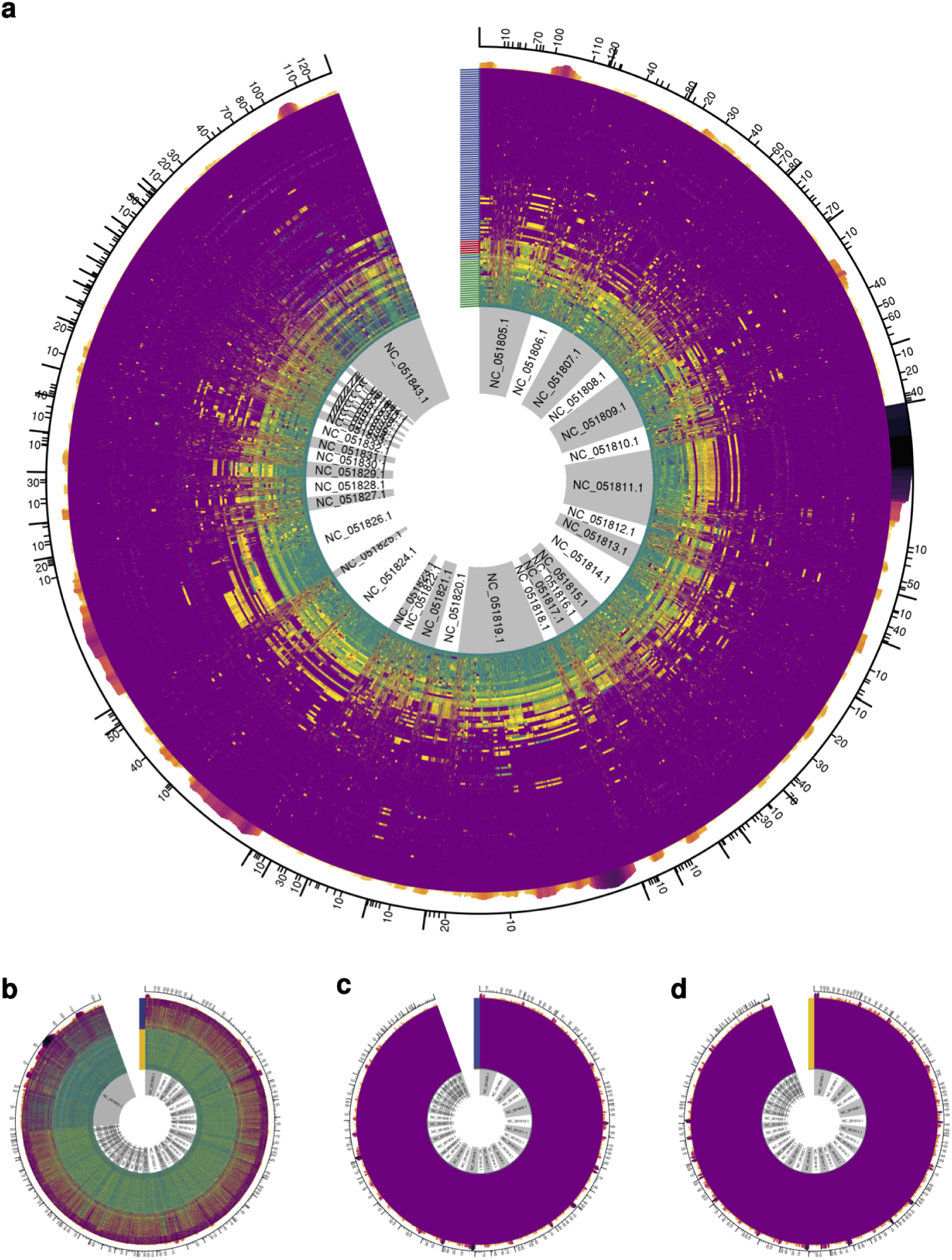
Polarised genomes demostrating gene flow in canids. Top 1% of markers with the highest diagnostic index are displayed. Tick mark colours correspond to taxon labels in Figure 1a. Barplot on top of the polarised genomes shows marker enrichment density (number of the most diagnostic markers per 1M bp) to site density (number of all variant sites per 1M bp). **a**, 86,434 diagnostic markers in Nearctic wild canids; coyotes (*n* = 23), red wolves (*n* = 6), and grey wolves (*n* = 78). High resolution iris plots for subsets Eurasia (**b**), jackalWolf (**c**), and jackalDog (**d**) are in the Supplementary Figures 3–5.

As wolves repopulate areas with high human densities, encounters with free-ranging dogs increase, leading to ongoing hybridisation. This issue is particularly acute in Western Eurasia, where dog admixture in wolves is higher than in East Asia and North America [10, 19, 21, 39]. The higher levels of hybridisation in Europe correlate with the region’s dense human population, as human activities, such as hunting, exacerbate this issue by disrupting wolf pack structures, making hybridisation more likely [21]. For example, wolf-dog hybrids are common in Italy [19], but in Fennoscandia, no wolf-dog hybrids were detected [40]. This variability underscores the need for targeted conservation strategies, including the development of reliable genetic markers to identify hybrids [11, 12, 41].

Due to the high admixture of coyote genomes in wolves from the Nearctic (Figure 2, Supplementary Figure 2a), we assessed gene flow between dogs and wolves concentrating on the Palearctic. The subset of the whole genome dataset included 675 individuals and 13,937,682 variant sites (Supplementary Table 1, subset Eurasia). Genome polarisation revealed distinct groups corresponding to the Palearctic wolves and dogs (Figure 2b, Supplementary Figure 3). The individuals with the rescaled hybrid index in the intermediate values between 0.4 and 0.75 included known F1 hybrids between wolves and dogs [40], dog hybrid breeds that represent backcrosses back to the dog gene pool [42], two free-ranging dogs from China (SAMN29845486, SAMN29845400 [42]), and four wolves. The four wolves with high proportion of dog admixture are from Israel (SAMN03366710) and three different regions in China (Himalayas: SAMN12565085, Shanxi: SAMN03168395, Inner Mongolia: SAMN03168393) (Supplementary Table 1) [43, 44].

Introgressions between wolves and dogs were difficult to identify, as Scandinavian wolves were largely homozygous and formed one side of the barrier to gene flow (Figure 2b, Supplementary Figure 3). In contrast, most Palearctic wolves were heterozygous at the 1% of markers with the highest diagnostic index, which masked potential dog introgressions. On the dog side, our results show that the F1 hybrids were all male, with wolf mothers (yellow horizontal lines changing to purple in X chromosome; Supplementary Figure 3). Wolf introgressions were observed in hybrid dog breeds, though most other dogs showed only small sections of wolf ancestry.

Some studies showed that golden jackals contributed to gene pool of dogs through hybridisation [45, 46]. Our results from 369 samples and 13,085,898 variant sites show no diagnostic markers originating from jackal genomic diversity in domestic dogs (Supplementary Figure 5). Similarly, diagnostic genomic variants distinguishing between 296 golden jackals and wolves based on a subset of 14,981,765 variant sites do not show geneflow into the wolf genome (Supplementary Figure 4).

## Wolf-dog hybrid breeds

The genomic diversity of domestic dogs has greatly diminished, as most of today’s dog breeds originated roughly 200 years ago from a small pool of founders. The strict enforcement of breed standards has further reduced genetic diversity and led to increased inbreeding [47]. Despite this reduction in genomic diversity, domestic dogs remain among the most phenotypically diverse mammals. This diversity, however, arises from relatively few key genomic changes rather than extensive genetic differences [48, 49].

The wolf-like phenotype has long been attractive to dog breeders, though early generation hybrids often present behavioural challenges that make them difficult to manage in domestic settings. Two dog breeds, the Czechoslovakian and Saarloos wolfdogs, resemble wolves and have recent wolf ancestry [22, 42, 50]. The Czechoslovakian Wolfdog was established in 1958 from a female Carpathian wolf and a German Shepherd male. To further enhance the breed’s characteristics, additional introductions of two male wolves and another female wolf into the breeding stock occurred between 1960 and 1983 [22, 23]. Similarly, the Saarloos Wolfdog traces its origin to a cross between a German Shepherd male and a female wolf from Siberia in 1935. Genetic studies consistently highlight the significant wolf ancestry in these breeds [22–24, 42, 51, 52].

On the other hand, breeds like the Northern Inuit Dog and Tamaskan were developed within the last 40 years from multiple other dog breeds, including the Alaskan Malamute, Siberian Husky, and German Shepherd, to achieve a wolf-like appearance with dog-like behaviour. No direct wolf involvement in their breeding is documented, and there is also no recent wolf admixture reported in the ancestral breeds [53]. However, the ancestral breeds Alaskan Malamute and Siberian Husky are ancient breeds closely associated with Paleolithic dogs from Siberia [54, 55]. Both carry the *MC1R* gene mutation p.R301C often used as a marker of wolf ancestry [49, 56]. In contrast, the German Shepherd is a modern breed of European origin with no reported wolf ancestry or ancient genetic markers.

Here, we identified the introgressed blocks in two hybrid dog breeds, namely the Czechoslovakian Wolfdog, and the Saarlos Wolfdog, and two dog breeds with wolf-like phenotype, the Northern Inuit Dog and the Tamaskan. We also include expected parental breeds of these breeds, the Alaskan Malamute, Siberian Husky and German Sheperd (Figure 1). The dataset is a subset with 162 samples and 17,268,030 variant sites. As expected, genomes of the hybrid dog breeds show close affinity to wolf genomes, but the mosaic of the introgressed blocks of wolf origin differs (Figure 3).

**Figure 3:**
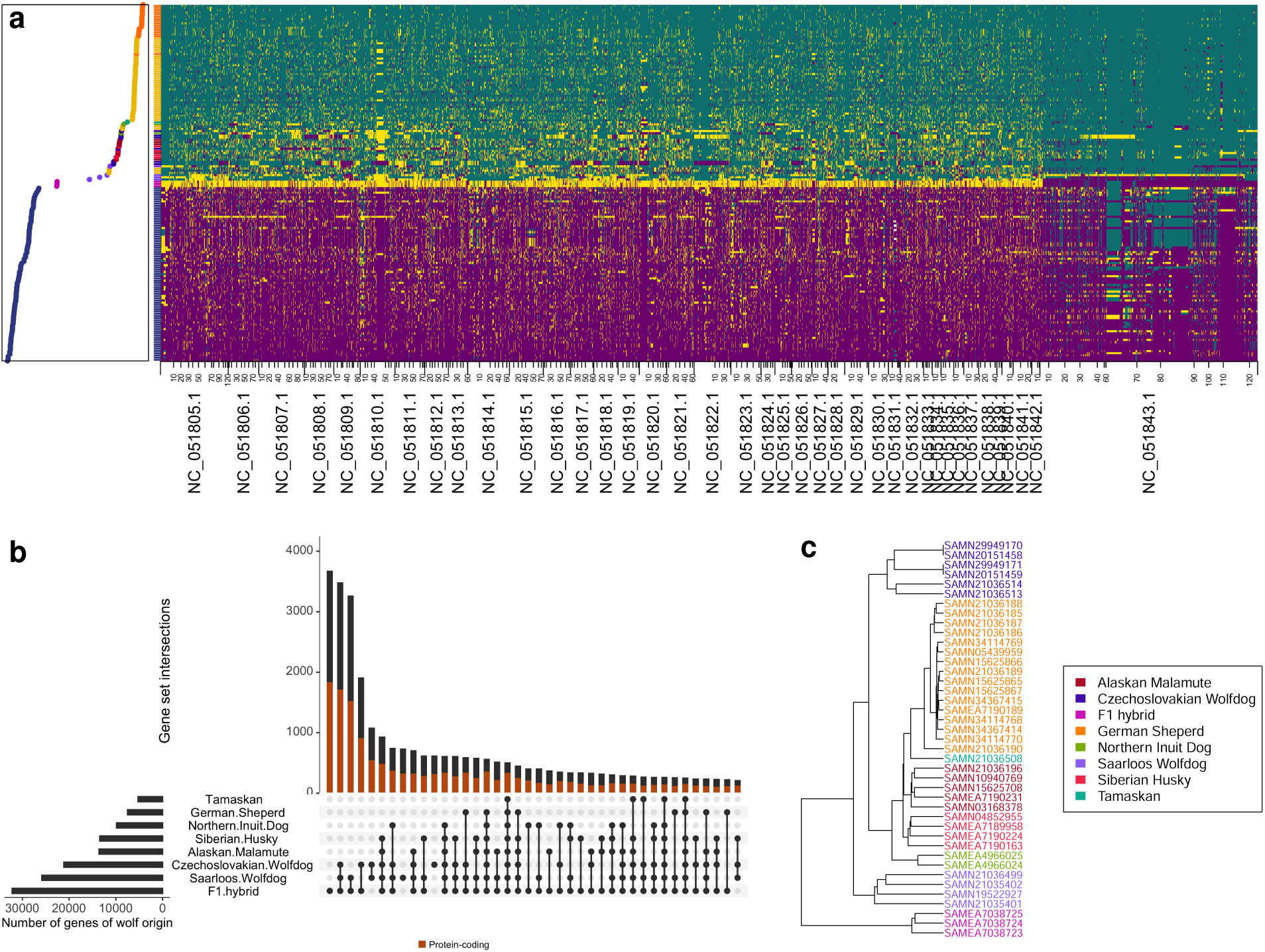
Genome admixture in modern hybrid dog breeds. **a**, Polarised genomes of wolves from Eurasia and modern dog breeds (Supplementary Table 1, subset dogHybrids). Top 1% of the most diagnostic markers is shown, with the genomic states being smoothed to the weighted mode of polarised states 125kb up- and down-stream from each site. Hybrid index to the left of the polarised genomes and the corresponding rug are colour-coded according to the scheme in Figure 1a. Purple – homozygous genotypes for the wolf allele, green – homozygous genotypes for the dog allele, yellow – heterozygous genotypes, white – missing data. **b**, Intersections of gene sets in dog breeds containing alleles from the wolf side of the barrier to gene flow. F1 hybrids are included as a control. **c**, Complete-linkage dendrogram of Manhattan distances between dog breed gene sets that contain alleles from the wolf side of the barrier to gene flow.

Saarlos Wolfdogs are most closely related to the wolves, which is largely attributable to their wolf-like X chromosome (Figure 3). Czechoslovakian Wolfdogs have large wolf-like genomic blocks across the X chromosome, but also on chromosomes 4, 5, 18, and 26. The introgressed blocks are smaller in the other breeds, and they share limited genomic content (Figure 3).

## Genome architecture of marker diagnosticity enrichment

We examined the chromosomal distribution of the top 1% of markers with the highest diagnostic index and found that their enrichment is non-random (Figure 2, Supplementary Figure 6). Marker enrichment density, defined as the ratio of high-diagnostic-index markers per 1 Mb to the total number of variant sites per 1 Mb, was consistently elevated near the beginnings of chromosomes. The strength and direction of correlation between marker enrichment density and chromosomal position varied across subsets. In the jackalDog and jackalWolf comparisons, most chromosomes showed a significant negative correlation with position (Spearman’s correlation with significance assessed via circular permutation: *p*_FDR_ ≤ 0.05). In the Eurasia subset, chromosomes 1, 16, 18, 19, 21, 25, and 36 were significantly negatively correlated with position. In the NAwild subset, chromosomes 7, 11, and 16 showed significant negative correlations, while chromosome 38 showed a positive correlation.

In contrast, the overall density of variant sites tended to increase toward chromosome ends, with a significant positive correlation with site position across most chromosomes and subsets (Supplementary Figure 6).

We found no consistent correlation between marker enrichment density and GC content. The only exceptions were chromosomes 5 and 31 in the NAwild subset, where enrichment density was significantly positively correlated with GC content (Supplementary Figure 6a).

## Discussion

One of the key strengths of our approach is the use of whole-genome sequencing, which avoids the dog-centric bias inherent in SNPchip-based studies [21]. SNPchips often rely on pre-selected markers that can skew results, especially when designed with a specific species in mind, such as domestic dogs. By contrast, whole-genome sequencing provides a more comprehensive and less biased view of genetic diversity. However, our method is not without its challenges. The variant site selection process tends to focus on the two most frequent alleles at each site, which, in an imbalanced dataset (Figure 1a), often includes a dog allele. Consequently, while our data is more robust and less biased than many other datasets, it is not flawless, as we may lose third and fourth alleles that could be important for understanding the genetic diversity of rarer taxa in our study.

### Ancestral polymorphism in dogs

The primary axis of genomic diversity among grey wolves, dogs, coyotes, red wolves, and jackals extends from jackals to Mexican wolves. Notably, the Syrian jackal exhibits the highest jump in hybrid index (Figure 1), yet there are no significant barriers to gene flow among these taxa. Instead, genomic admixture gradually increases from coyotes to Mexican wolves. A key area of interest lies in the upper end of the genomic admixture continuum. Mexican wolves cluster closely with wolves from the Tibetan Plateau and Greenland, as well as with a group comprising Pleistocene wolves and dogs. Bergström et al. [27] demonstrated that dogs are more closely related to Pleistocene wolves from eastern Eurasia than to contemporary wolves, and phylogenetic investigation of mitochondrial genetic diversity [34] supports our whole-genome finding that the wolves in New Mexico represent ancient wolf genetic diversity now extinct across most of the wolf distribution. Our study builds on these findings by incorporating a broader sample set and leveraging whole-genome sequencing, revealing that ancient genetic diversity has been preserved in wolves from isolated regions such as New Mexico, Greenland, and the Tibetan Plateau (Figure 1). Outside of these three regions, domesticated dogs represent the primary reservoir of ancient wolf genetic diversity, a diversity that has been largely lost in wolf populations globally (Figure 1, Supplementary Figure 1).

The observed discrepancy between the retained historical diversity in dogs and the divergence of wolves since the Pleistocene may result from a relaxation of selective pressures in dogs. During domestication, humans imposed selective pressures on a limited set of genes related to desired phenotypic traits [48, 49, 55, 57]. This selective breeding focused on specific characteristics while leaving the majority of the dog genome free from natural selection pressures, facilitated by the protected environment of human settlements that provided consistent food resources.

### Canid hybridisation in the Nearctic

The three Nearctic canids, coyotes, red wolves, and grey wolves, exhibit two barriers to gene flow that do not fully align with the species classifications in the NCBI database (Figure 2). All individuals with a hybrid index below 0.75 are identified as coyotes, while all individuals with a hybrid index above 0.8, except for one red wolf presumably from Texas, USA [16], are classified as grey wolves. The greatest taxonomic diversity was observed in the group with whole-genome hybrid indices between 0.75 and 0.8, which included coyotes from the US Midwest, Missouri, and Alabama, as well as red wolves and Eastern wolves (*Canis lupus lycaon*) [13, 58–60]. Multiple studies have explored the systematics of Nearctic canids, particularly the taxonomic status of the red wolf and Eastern wolf, and the known hybridisation between coyotes and wolves. Our data provide a one-dimensional perspective on the genetic diversity of Nearctic canids, clearly indicating weak, semi-permeable barriers to gene flow among these taxa. These data-driven sample assignments to specific taxonomic groups offer a clear and traceable guide for understanding sample membership. However, the genomic continuity among Nearctic canids poses challenges to traditional species concepts, revealing a gradient of genetic variation that complicates species delineation.

### Wolf-dog hybridisation

Our findings demonstrate that gene flow between Palearctic wolves and domestic dogs, while present, is less pronounced compared to the extensive hybridisation between coyotes and wolves in the Nearctic (Figure 2). Genome-wide analyses revealed that most wolves and dogs formed distinct clusters, with only a few individuals showing intermediate hybrid indices (Figure 2).

Having dialed the genome polarisation to a subset of individuals expected to form a single barrier to geneflow (compare Figures 1 and 2), and filtered the sites to obtain the most diagnostic markers for that specific barrier, one would expect F1 hybrids to have a rescaled hybrid index of 0.5, as they inherit a set of chromosomes from each parent. The hybrid index of the known F1 hybrids in this study is less than 0.5. This is due to the fact that all included F1s were male descendants from a female wolf, having inherited a wolf-like X chromosome (Figure 3). This discrepancy is important, as the X chromosome carries the highest proportion of diagnostic markers across the barriers studied herein. The increased frequency of diagnostic markers on the X chromosome is likely due to its mode of inheritance; the X chromosome is maternally inherited in males and remains unpaired, leading to reduced recombination and preservation of lineage-specific alleles, making it a more frequent source of diagnostic markers [61].

Importantly, while four wolves exhibited genomic evidence of hybridisation with dogs, the over-all extent of hybridisation in the Palearctic appears limited compared to the Nearctic system. The risk of hybridisation remains a concern, particularly in densely populated regions, where human activities, such as hunting and intensive agriculture, disrupt wolf packs and increase encounters with free-ranging dogs [19, 21]. Although we identified early generation wolf-dog hybrids in Israel and China, other countries with samples of wolves with relatively high hybrid index indicating gene flow between dogs and wolves are Croatia, India, Iran, Italy, Kyrgyzstan, Saudi Arabia, South Korea, Spain and Syria (Supplementary Table 1). Low hybrid indices in wolves from Norway and Sweden are to be expected, as the wolves in Scandinavian peninsula recovered from a very low number of founders in the 20th century, with limited immigration [37, 40, 62].

### Introgressions in hybrid dog breeds

Hybrid dog breeds serve as valuable evolutionary experiment for studying wolf introgressions in grey wolves. With documented wolf ancestry, the Czechoslovakian and Saarloos Wolfdogs illustrate how recombination and backcrossing to dogs reshape the genomic composition of these breeds. Our results indicate that the Saarloos Wolfdog retains the highest proportion of wolf-like alleles, with significant wolf ancestry concentrated on the X chromosome and large blocks (over 20 Mb) on chromosomes 1 and 5 (Figure 3a). Smaller wolf-derived blocks are scattered across the genome, mostly in a heterozygous state. The wolf-like alleles in Saarloos Wolfdogs are located in approximately 1,800 protein-coding genes shared with F1 hybrids, but not present in other dog breeds (Figure 3).

In contrast, the Czechoslovakian Wolfdog displays several blocks with wolf-like alleles in homozygous state across its genome (Figure 3). These larger genomic blocks reflect the more recent hybridisation with wolves, around 50 years ago, compared to the nearly 90 years since the first wolf-dog cross in Saarloos Wolfdogs [22, 23, 50]. The Czechoslovakian Wolfdog has around 900 unique protein-coding genes with wolf-like alleles, with an additional 2,000 shared with other hybrids (Figure 3).

As expected, the Northern Inuit Dog and Tamaskan, which were bred for wolf-like appearance but lack direct wolf ancestry, carried minimal wolf-like genomic content. Both breeds contained fewer wolf-like alleles than their ancestral breeds, the Alaskan Malamute and Siberian Husky, which themselves retain limited ancient variation (Figure 3). The number of protein-coding genes with wolf-like alleles was more than an order of magnitude lower in Northern Inuit Dogs and Tamaskans compared to the hybrid wolfdogs. Across all four wolf-like breeds, fewer than 200 protein-coding genes were shared, underscoring the limited overlap and supporting the known ancestry of these phenotypically wolf-like but genetically dog-derived breeds.

These findings highlight that domestication, focused on selecting specific traits, affects only a small fraction of the genome [48]. The persistence of wolf-like alleles in hybrid breeds underscores how selective breeding and backcrossing can maintain or lose ancient genetic diversity, depending on the breed’s history and the timing of hybridisation events. The absence of significant wolf-like introgressions in the German Shepherd support this, reinforcing the German Sheperd’s distinct status as a domestic dog. It is important to remember though that the genomic content of dogs positions them close to ancient wolves, emphasising their role as genetic reservoirs of ancestral diversity.

### Chromosomal structure and diagnostic marker enrichment

The strong enrichment of diagnostic markers at the beginnings of chromosomes points to elevated marker diagnosticity in pericentromeric regions (Supplementary Figure 6). This pattern aligns with the architecture of the canid genome, where nearly all autosomes are acrocentric, placing the centromere near the physical start of the chromosome. Consequently, the enrichment we observe corresponds spatially to these pericentromeric regions. The exception is chromosome 27, which appears inverted in the current reference assembly, consistent with the observed shift in enrichment toward the chromosomal end.

Recombination rates are typically reduced around centromeres and elevated toward telomeres. As GC content correlates with recombination in mammals [63], our finding that marker enrichment density does not consistently correlate with GC content suggests that the diagnosticity signal is not solely driven by local recombination rate. Instead, the accumulation of diagnostic markers in pericentromeric regions may reflect reduced recombination in combination with increased lineage sorting. Additionally, these regions may be subject to reinforcement, where selection acts more effectively against introgressed alleles due to limited recombination, thereby maintaining stronger species boundaries.

## Conclusions

While it is well established that grey wolves hybridise with sympatric canids, most notably with coyotes and red wolves in the Nearctic and with domestic dogs in the Palearctic, our findings reveal more than just the persistence of gene flow. Through genome polarisation across five canid taxa, we uncover a structured pattern of marker diagnosticity that reflects deeper evolutionary dynamics. Diagnostic markers are not evenly distributed across the genome, but are disproportionately concentrated in pericentromeric regions, areas of low recombination that appear to reinforce species boundaries. This pattern suggests that these regions act as genomic barriers, where reduced recombination and potentially reinforcement maintain lineage-specific alleles and restrict introgression.

We further show that ancient wolf diversity persists only in a few isolated wolf populations, such as those in New Mexico, Greenland, and the Tibetan Plateau, and, crucially, in domesticated dogs. Dogs represent a genomic arch for ancient wolf variation, preserving lineages otherwise lost in most wolf populations. Together, these results highlight that the architecture of the genome plays a key role in maintaining species integrity. The accumulation of diagnostic markers in pericentromeric regions suggests that reinforcement in low-recombination regions may be a general mechanism for preserving species boundaries across canids.

## Methods

### Sample selection

We selected a set of publicly available wolf and dog genomes sampled from their entire distribution range (Supplementary Table 1). For dogs, our focus was primarily on free-ranging dogs, particularly from areas where wolves naturally occur, supplemented with various pedigree and hybrid dog breeds. Additionally, we included data for both coyotes and red wolves to explore their role in hybridisation with the wolves in North America, historical sequences of Pleistocene wolves to investigate genomic divergence over time, and data from golden jackals to investigate ancestral polymorphisms in dogs.

BioProjects were filtered using NCBI run selector to remove irrelevant taxa, accessions smaller than 1 GB in size, and any non-genome accessions. Additionally, we excluded samples with repeated downloading and mapping issues. This resulted in a selection of 1313 canid genomic short read archives for 979 unique BioSamples (Supplementary Table 1).

Due to lack of wolves from the Carpathians, which is the region of origin of ancestral wolves for the Czechoslovakian Wolfdogs, we enriched the dataset with a grey wolf from Slovenský raj, Slovakia. The sample was a road kill from 2016 deposited in the Genetic Bank of the Institute of Vertebrate Biology. The sample was commercially sequenced on NovaSeq X Plus Series using paired-end sequencing.

### Mapping and variant calling

We mapped the reads onto the Labrador Retriever genome assembly ROS Cfam 1.0 (NCBI Genome Accession: GCA 014441545.1), a haploid reference constructed from PacBio sequencing of a single male [64].

We downloaded the reads for each sample using SRAtoolkit [65], and mapped them to the indexed reference genome using bwamem [66]. Read annotation, format conversion, sorting, duplicate marking, and read indexing were performed using Picard and Samtools toolkits [67, 68]. We then used these pre-processed bam files to carry out variant calling using bcftools mpileup and call functions [67].

Due to prevalent read overmerging, reads mapping to the unmapped contigs of the reference genome were excluded from subsequent analyses.

### Genome polarisation

Genome polarisation assumes neither marker diagnosticity nor sample affinity to either side of a barrier to geneflow. Rather, it provides estimates of these, inferred from the data through marker state associations across the whole genome [8]. This is particularly useful when studying wolf hybridisation, because genome polarisation avoids many pitfalls associated with the topic in earlier studies, such as insufficiently representative samples, incorrect or incomplete assignment of samples to specific groups, and bias of pre-selecting genetic markers assumed to be diagnostic [24, 69]. Genome polarisation allows inclusion of many samples, bringing representativeness to a higher level, does not require one to know what groups the samples belong to, and can include all variant markers across many genomes.

Before polarising the dataset or its subsets, single nucleotide variants (SNVs) called in VCF files were converted into *diem* marker format using vcf2diem [8]. At each SNV site, the two most frequent alleles were identified and encoded into ‘0’ and ‘2’ for homozygotes, ‘1’ for heterozygotes, and **‘ ’** for indels, missing data or genotypes with other alternative alleles. This step also served as a quality control and data filtration measure, omitting sites lacking homozygotes for both most frequent alleles.

Subsequently, we polarised the dataset using the *diem* algorithm for genome polarisation implemented in the R package *diemr* 1.4.3 [8]. The *diem* algorithm first randomly re-assigns which homozygote is encoded ‘0’ and ‘2’ for each marker, and then infers the side of a barrier to which each allele belongs using expectation maximisation. This method allowed us to efficiently handle large quantities of markers (in the tens of millions), and to compute the diagnosticity of each marker from the data without any reliance on priors, such as sample purity and barrier side association, or usage of markers previously asserted to be diagnostic [8, 70]. For subsequent analyses and plotting, we used the output diagnostic index of markers for filtering and marker selection.

We ran genome polarisation with different sets of individuals to assess different axes of gene flow. As genome polarisation finds the strongest barrier to gene flow in a given dataset, it can be applied to differentiate between distinct groups of genomic variation. To identify groups of samples of interest, we used sample hybrid indices (*h*, proportion of the genome belonging to one side of the barrier rescaled to the interval [0, 1]) calculated either from the whole dataset or from a fraction of the markers with the highest diagnostic index. In combination with sample labelling in the database, we established a *h* threshold for splitting the dataset. We used this genome-state-driven approach to eliminate the influence of inconsistent or incorrect sample labelling in the databases to identify samples with admixed genomes.

When analysing subsets of samples, variant sites were re-filtered using vcf2diem, requiring that each marker include a minimum number of homozygous individuals for both of the two most frequent alleles. This threshold was increased to reduce the influence of outliers or inbred individuals on genome polarisation. For the jackalDog and jackalWolf subsets, the minimum was set to two; for all other subsets, to five. Reformatting the data for each subset also allowed the selection of a different allele pair than in the full dataset, capturing variants where alternative alleles were more frequent in the subset than globally.

Genome polarisation randomly assigns either side of the barrier as being associated with low values of the hybrid index. This lack of directionality can lead to the subset being ‘flipped’ compared to the full dataset [8]. In such cases, we enforced the directionality of the detected barriers in the subsets to correspond to their directionality in the analysis of the full dataset following recommendations in [8].

To investigate introgressions, we smoothed marker states across the diagnostic markers to remove noise from standing variation orthogonal to the barrier signal [70, 71]. We used a sliding-window approach, were the genomic state at each diagnostic marker was updated as a weighted mode of the genomic state for a window that included 0.25M sites centered on the diagnostic marker. We used truncated Laplace probability densities in such a way that the curve integrated to 0.95, and the scale parameter was equal to 41726.03. The probability densities for the physical distance (in bp) from the marker being smoothed were then scaled proportionally so that the smoothed marker had weight equal to 10*/*19. Note that all distances in the smoothing were physical distances along the reference genome, therefore avoiding bias caused by different density of variant sites along chromosomes.

Genome polarisation followed by diagnostic marker genomic state smoothing facilitates identification of introgressed blocks without the need for pure populations and perfectly diagnostic markers. This is an advantage with respect to identification of runs of homozygocity (ROH), as the introgressed blocks will be highlighted also in heterozygous states.

### Diagnostic marker enrichment density

We quantified the enrichment of diagnostic markers along chromosomes by comparing their local density to the overall density of SNVs. For each SNV, we counted the number of SNVs within a 1 Mb window. We then identified the top 1% of diagnostic markers based on their diagnostic index and calculated the number of these highly diagnostic markers within the same 1 Mb window. The enrichment fold-change at each site was defined as the ratio of diagnostic marker density to total SNV density.

To assess structural variation in diagnostic marker enrichment, we analysed its relationship with chromosomal position, local SNV density, and GC content. GC content serves as a proxy for recombination rate, which influences the genomic distribution of variants [63]. We tested for significant associations using Spearman’s correlation, estimating a null distribution via circular permutations.

Circular permutations provide a robust framework for detecting structural variation along chromosomes, accounting for autocorrelation in genetic variation [71]. To generate the null distribution, we applied random circular shifts of up to 20% of chromosome length, preserving relative spacing while disrupting direct positional associations. We then calculated the significance of observed correlations by comparing them to the permuted null distribution, applying false discovery rate (FDR) correction to adjust for multiple testing.

## Acknowledgements

Samples were provided by the Genetic Bank of the Institute of Vertebrate Biology, Czech Academy of Sciences. This study was supported by the Czech Academy of Sciences, Institute of Vertebrate Biology (RVO:68081766). Authors thank the RECETOX Research Infrastructure (No LM2023069) financed by the Ministry of Education, Youth and Sports for supportive background. Computational resources were provided by the e-INFRA CZ project (ID:90254), supported by the Ministry of Education, Youth and Sports of the Czech Republic.

## Author Contributions

Conceptualisation, N.M.; Design, F.J, S.J.E.B and N.M.; Investigation, F.J.; Methodology, F.J. and N.M.; Formal Analysis, F.J., S.J.E.B., M.H., and N.M.; Writing – Original Draft, F.J., S.J.E.B., M.H., and N.M.; Writing – Review & Editing, F.J., S.J.E.B., M.H., and N.M.

## Data Accessibility

Raw sequence reads are deposited in the Sequence Read Archive (BioProject PRJNA1170880). Additional datasets used in the study are listed in Supplemetary Table 1.

## Supplementary Information

**Supplementary Figure 1:**
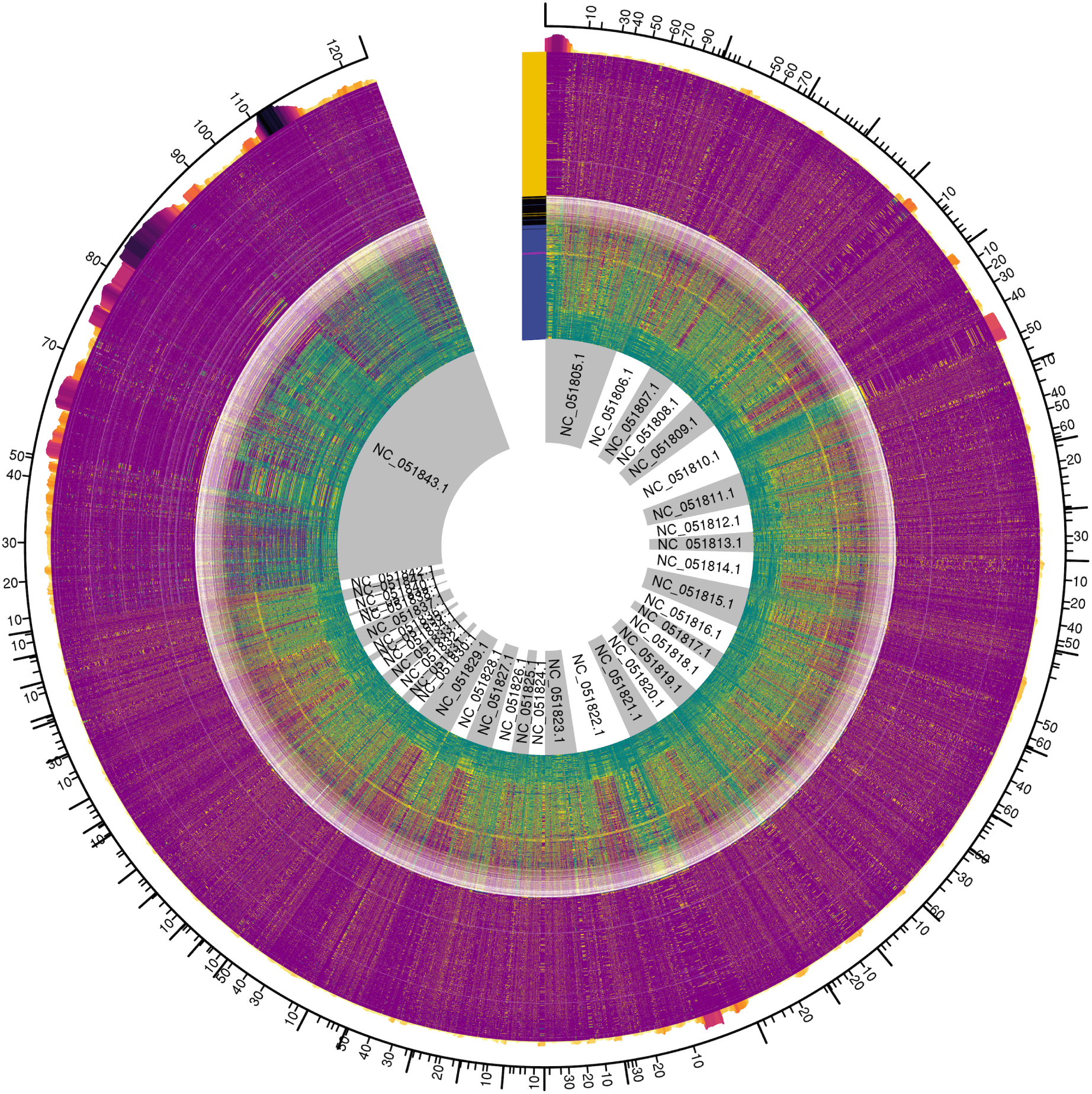
High resolution polarised genomes with focus on a barrier to gene flow between grey wolves (*n* = 364), including 71 ancient wolves and dogs (*n* = 379). Top 1% of markers with the highest diagnostic index are displayed (139,822 markers). Tick mark colours correspond to those in Figure 1a. Barplot on top of the polarised genomes shows ratio of marker density (number of the most diagnostic markers per 1M bp) to site density (number of all variant sites per 1M bp).

**Supplementary Figure 2:**
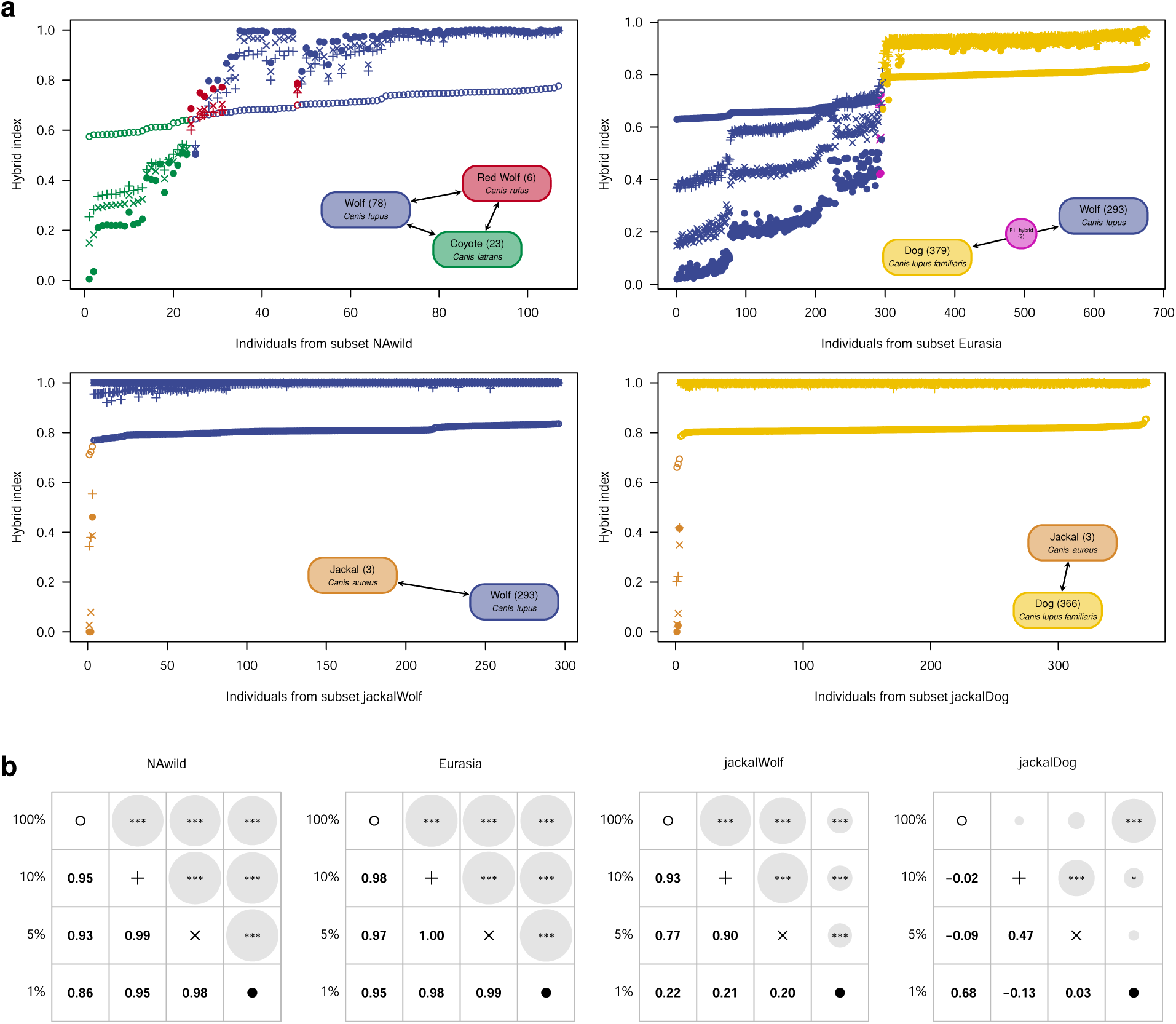
Reproducibility of the hybrid indices across canid barriers to gene flow, respective to the sampled proportion of the genome. **a**, Sorted hybrid indices for canid subsets (Supplementary Table 1) corresponding to separate barriers to gene flow (Figure 1b). The stability of hybrid indices is demonstrated across levels of diagnosticity, with increasing resolution at the top 10% (+), 5% (*×*), and 1% (*•*) of the most diagnostic SNVs compared to the whole-genome hybrid indices (*◦*). Insets display colour legends and sample sizes in taxa included in the respective subsets. **b**, Pairwise Spearman’s correlation coefficients (below diagonal) and FDR-adjusted significance values (above diagonal; * indicates *p ≤* 0.05, ** *p ≤* 0.01, and *** *p <* 0.001) for hybrid indices displayed in panel a.

**Supplementary Figure 3:**
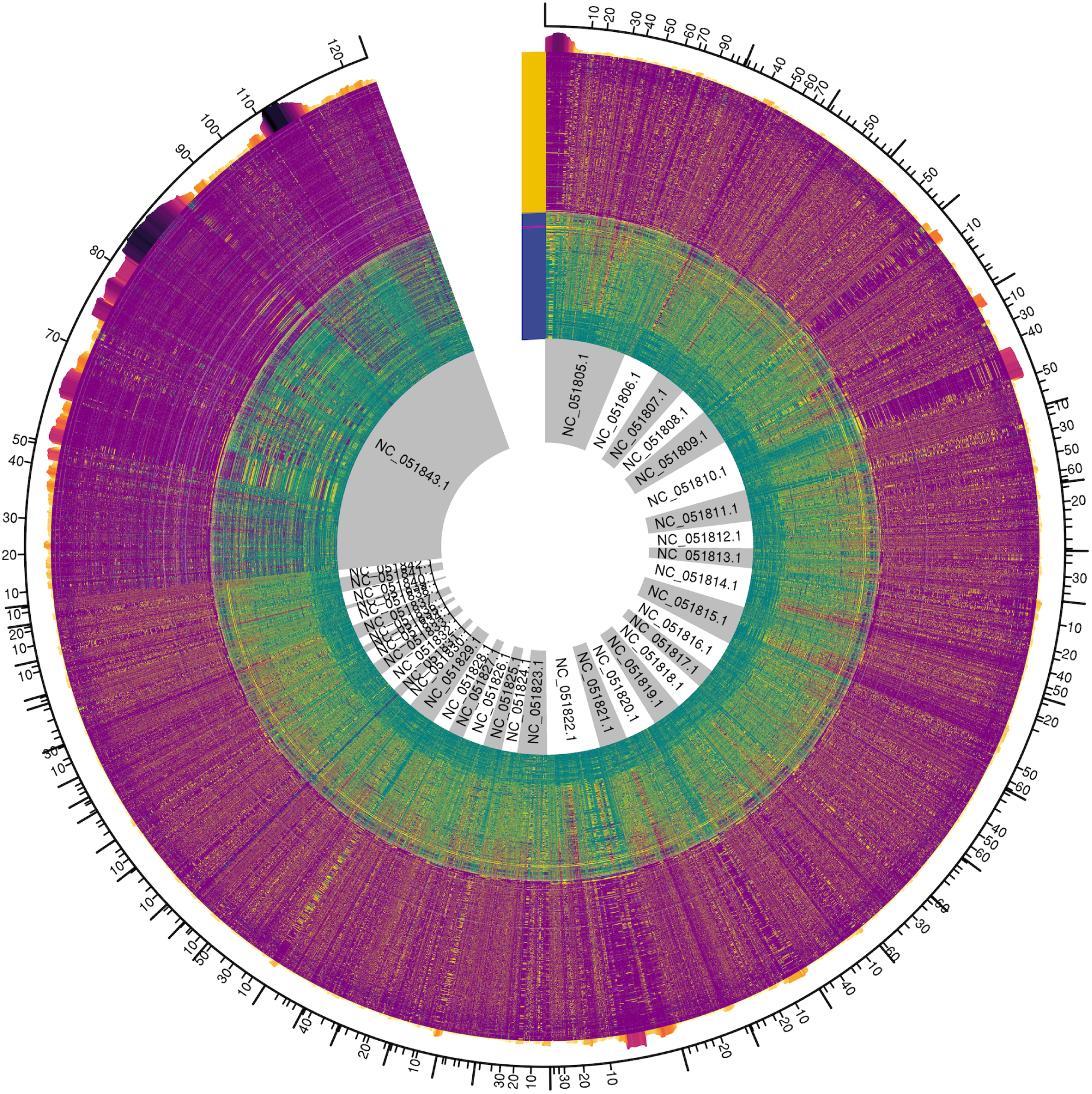
High resolution polarised genomes with focus on a barrier to gene flow between grey wolves from Eurasia (*n* = 293) and dogs (*n* = 382). Top 1% of markers with the highest diagnostic index are displayed (139,377 markers). Tick mark colours correspond to those in Figure 1a. Barplot on top of the polarised genomes shows ratio of marker density (number of the most diagnostic markers per 1 Mb) to site density (number of all variant sites per 1 Mb).

**Supplementary Figure 4:**
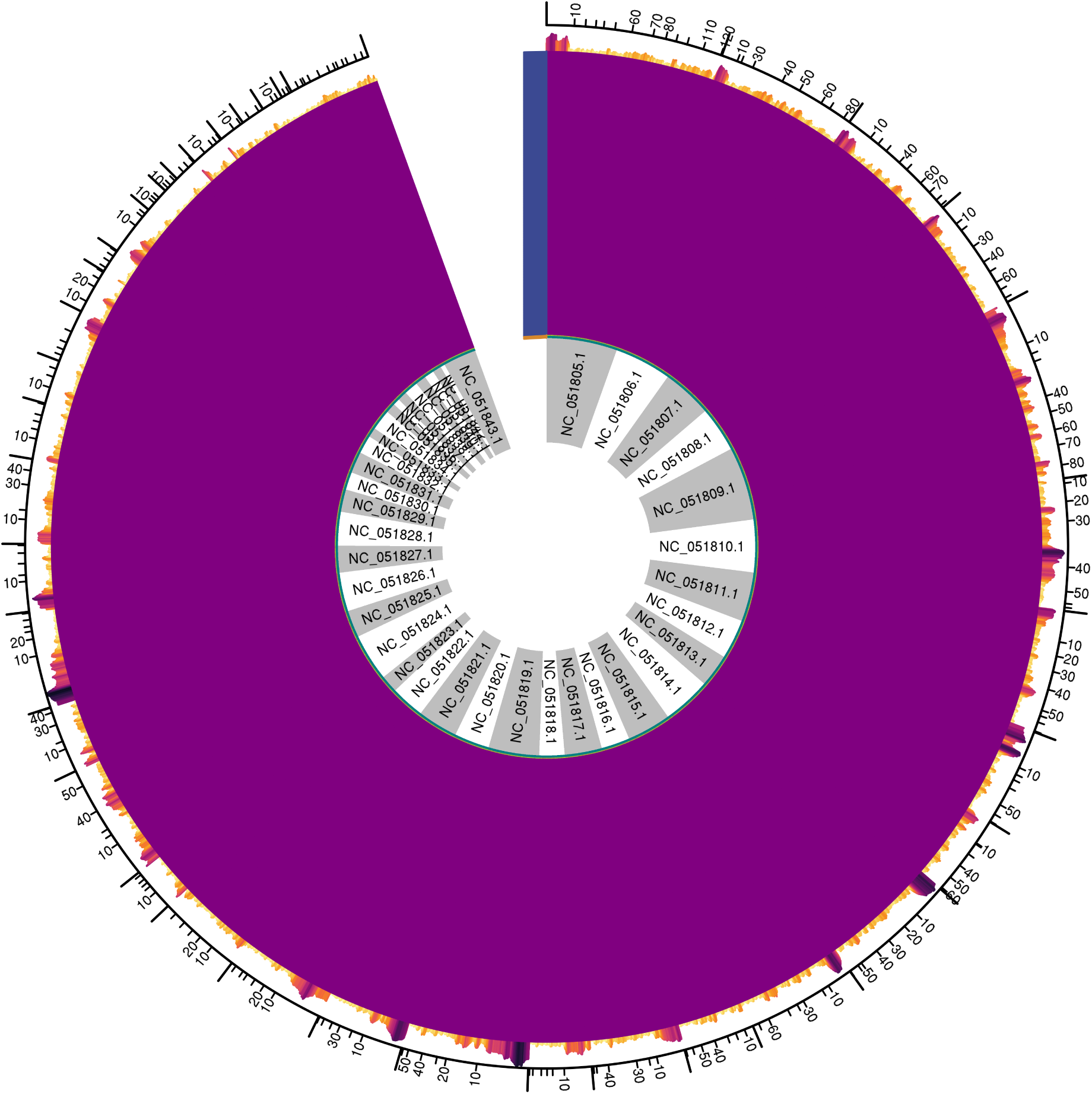
High resolution polarised genomes with focus on a barrier to gene flow between golden jackals (*n* = 3) and grey wolves from Eurasia (*n* = 293). Top 1% of markers with the highest diagnostic index are displayed (242,405 markers). Tick mark colours correspond to those in Figure 1a. Barplot on top of the polarised genomes shows ratio of marker density (number of the most diagnostic markers per 1 Mb) to site density (number of all variant sites per 1 Mb).

**Supplementary Figure 5:**
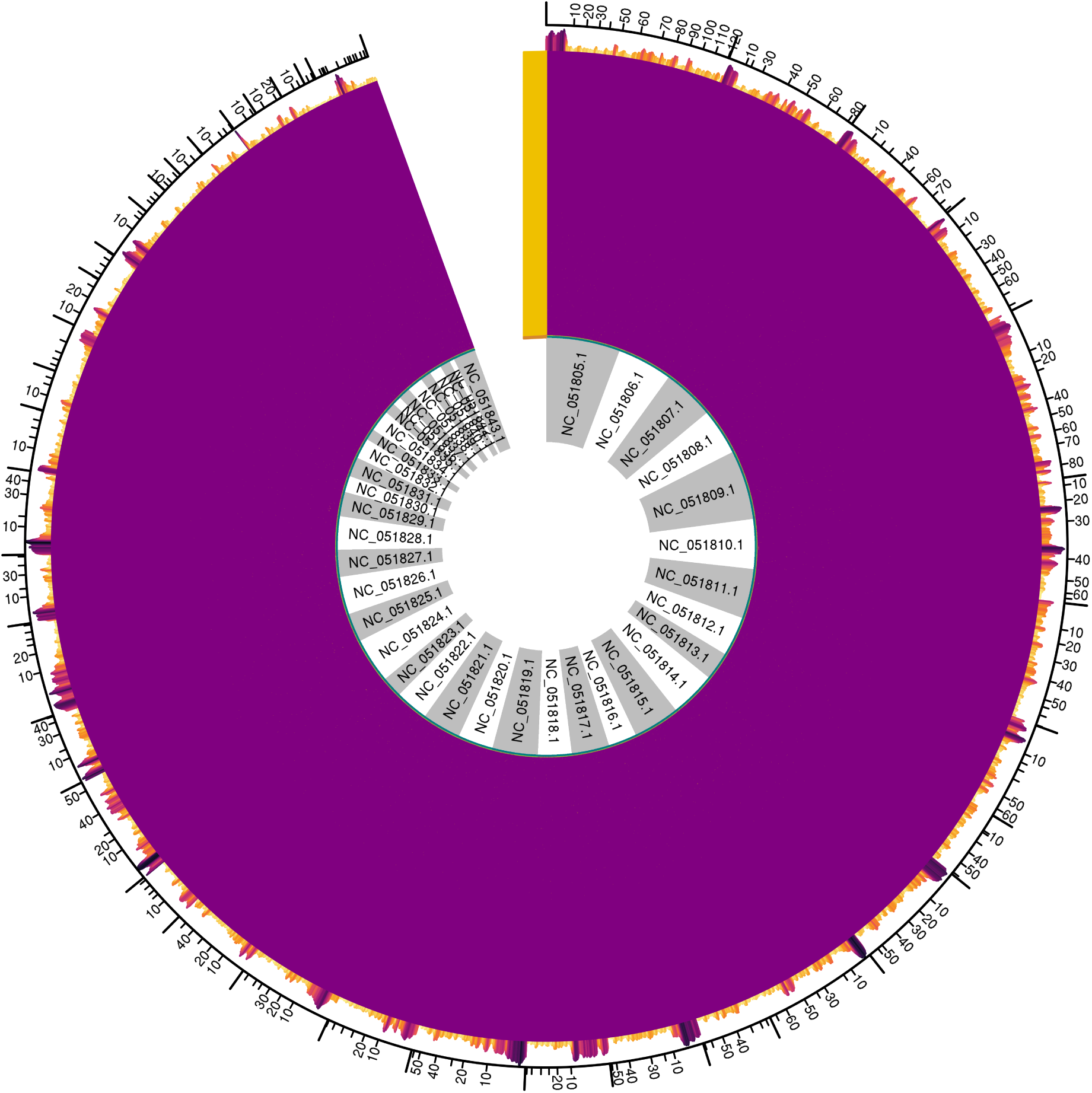
High resolution polarised genomes with focus on a barrier to gene flow between golden jackals (*n* = 3) and dogs (*n* = 366). Top 1% of markers with the highest diagnostic index are displayed (210,862 markers). Tick mark colours correspond to those in Figure 1a. Barplot on top of the polarised genomes shows ratio of marker density (number of the most diagnostic markers per 1 Mb) to site density (number of all variant sites per 1 Mb).

**Supplementary Figure 6:**
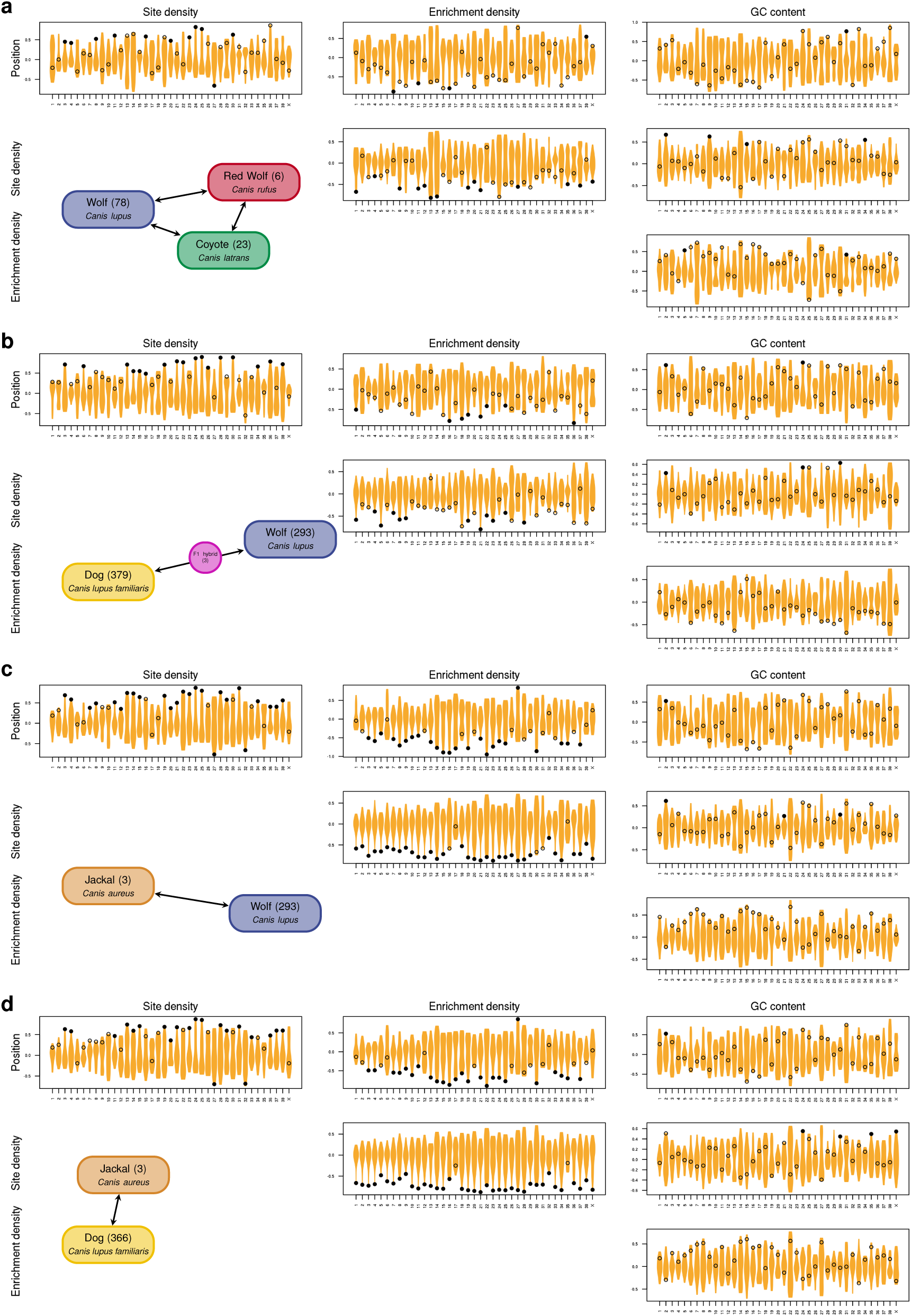
Structural variation along canid chromosomes comparing site position, variant site density, marker enrichment density (ratio of marker density (number of the most diagnostic markers per 1 Mb) to site density) and GC content, each in a 1 Mb window centered at the site position. Violin plots show pairwise Spearman’s correlation coefficients from circular permutations along each chromosome. Measured values are depicted as black circles that are filled when the measured correlation was significantly different from random (two-sided *p* values adjusted for multiple testing with FDR). Schemes of included taxa show number of genomes considered for the subsets. **a**, NAwild, **b**, Eurasia, **c**, jackalWolf, **d**, jackalDog.

Supplementary Table 1: Canid sample metadata and accession numbers. Table included as a separate, machine-readable file.

